# Imaging interorganelle phospholipid transport by extended synaptotagmins using bioorthogonally tagged lipids

**DOI:** 10.1101/2023.02.01.526230

**Authors:** Dongjun Liang, Lin Luan, Jeremy M. Baskin

## Abstract

The proper distribution of lipids within organelle membranes requires rapid, interorganelle lipid transport, much of which occurs at membrane contact sites and is mediated by lipid transfer proteins (LTPs). Our current understanding of LTP mechanism and function is based largely on structural studies and in vitro reconstitution. Existing cellular assays for LTP function use indirect readouts, and it remains an open question as to whether substrate specificity and transport kinetics established in vitro are similar in cellular settings. Here, we harness bioorthogonal chemistry to develop tools for direct visualization of interorganelle transport of phospholipids between the plasma membrane (PM) and the endoplasmic reticulum (ER). Unnatural fluorescent phospholipid analogs generated by the transphosphatidylation activity of phospholipase D (PLD) at the PM are rapidly transported to the ER dependent in large part upon extended synaptotagmins (E-Syts), a family of LTPs at ER-PM contact sites. Ectopic expression of an artificial E-Syt-based tether at ER-mitochondria contact sites results in fluorescent phospholipid accumulation in mitochondria. Finally, in vitro reconstitution assays demonstrate that the fluorescent lipids are bona fide E-Syt substrates. Thus, fluorescent lipids generated in situ via PLD activity and bioorthogonal chemical tagging can enable direct visualization of the activity of LTPs that mediate bulk phospholipid transport at ER-PM contact sites.

## INTRODUCTION

Every organelle has a distinct membrane lipidome, whose composition results from complex regulation of both lipid metabolism and intracellular transport. Recent studies have identified numerous non-vesicular pathways as primary players in controlling interorganelle lipid transport^1^. These pathways typically operate at membrane contact sites (MCS), which are sites of close apposition (< 30 nm) between membranes of different organelles, via lipid transfer proteins (LTPs) that can act variably as shuttles, bridges, or tubes^2^.

The physiological importance of interorganelle lipid transport is underscored by numerous diseases that can result from aberrant expression or mutation of LTPs, including neurodegenerative diseases^3–8^, lysosomal storage disorders^9,10^, and cancer^11–17^. Yet the ensemble of LTPs is exceedingly complex and challenging to study. It involves many players whose lipid specificities, transport kinetics, regulatory mechanisms, and extent of coupling to metabolic enzymes, scramblases, and/or flippases are in many cases still unknown^18^. These parameters can sometimes be determined for individual LTPs by in vitro reconstitution of lipid transport using purified proteins and liposomes of defined lipid composition. But, reminiscent of the vesicle fusion machinery, the in vitro specificities and kinetics of LTPs may not match those in a native cellular environment, and further, transport in cells is carried out by many LTPs acting in parallel.

Current approaches for monitoring intracellular lipid transport are indirect. Classically, pulse-chase labeling with radiolabeled lipid precursors or lipids, followed by *ex vivo* detection of resultant lipids (sometimes with an organelle fractionation step), can provide information on kinetics of lipid metabolism and interorganelle transport^19,20^. However, multiple redundant metabolic steps can confound analysis, and when used, fractionation typically affords only a partial enrichment. Genetically encoded fluorescent biosensors are powerful tools for reporting on lipid localizations in live cells and in real time^21,22^. However, they are more useful for yes/no decisions on whether a particular lipid is present in a membrane above a threshold amount, or how quickly its levels decay in a particular membrane. These biosensors are less well suited for quantifying the kinetics of dynamic lipid transport in between membranes because their relative affinities for the same lipid can differ in different membrane environments and their overall localizations result from a complex equilibrium of different affinities.

A recent method using a genetically encoded, organelle-tethered microbial enzyme that labels unsaturated phospholipid tails with a small tag detectable by mass spectrometry can enable non-radioactive tracking of changes in lipid localizations. This method, termed METALIC, uses heterologous expression of a bacterial cyclopropane fatty acid phospholipid synthase (CFAse) to tag unsaturated phospholipids. In conjunction with ER-localized PE methyltransferases (PEMTs) and metabolic labeling with stable isotope-labeled forms of a precursor for the mass tags installed by both CFase and PEMTs, METALIC can enable monitoring of bulk phospholipid transport between mitochondria and the ER in yeast.^23^ This powerful approach stands poised to reveal the kinetics of lipid transport mechanisms between different organelle contact sites, with numerous beneficial attributes, including monitoring near-native lipids and portability to different organelle contact sites and cell types. Yet, deconvolution of contributions from native lipids, particularly in mammalian cells, requires specialized mass spectrometry instrumentation and expertise.

Ideally, to trace interorganelle lipid transport directly in live cells, one would want a fluorescent lipid analog that is an LTP substrate, meaning that its changes in localization would directly report on LTP activity. The challenges to this approach are at least two-fold. First, fluorescent lipids differ substantially from native lipids and may have intrinsic localization preferences and may not be accepted by LTPs. Second, when a fluorescent lipid is added to cells exogenously, it is hard to specify a precise and desirable “start” position, which would, for assessing LTP function, preferably be a cytosol-facing leaflet. To surmount these issues, we envisioned building fluorescent lipids *in situ* using endogenous lipid biosynthetic enzymes operating at defined membrane locations, monitoring their subsequent redistribution by time-lapse microscopy, and using genetic perturbation by loss- and gain-of-function to establish the LTPs responsible for their trafficking. Then the fluorescent lipid would constitute a useful tracer for the identified step(s) in the intracellular lipid transport network.

In this study, we develop such a toolset for imaging lipid transport at ER-PM contact sites in large part by the extended synaptotagmin (E-Syt) family of LTP shuttles. ER-PM contact sites support many important physiological functions, including regulation of Ca^2+^ uptake and signaling^24^, export of phosphatidylserine^25^, cholesterol^20^, and other lipids from the ER to the PM^19,26,27^, and maintenance of the phosphoinositide cycle^28^, which regulates exo/endocytosis, actin dynamics, ion channel activity, and second messenger signaling. E-Syts (tricalbins in yeast) act at ER-PM contact sites to both tether the two organelle membranes in close proximity and also, via their central hydrophobic synaptotagmin-like mitochondrial-lipid-binding protein (SMP) domain, mediate interorganelle transport of glycerolipids^29^. Though they are only one of several families that can mediate lipid transfer at ER-PM contact sites, extensive in vitro and cell-based studies have established that they have a broad glycerolipid substrate preference^30^, suggesting that they may accommodate lipid analogs with unnatural head groups.

Here, we exploit bioorthogonal chemistry-powered tools using unnatural fluorescent phospholipids for visualizing LTP activity at ER-PM contact sites in live cells mediated in large part by E-Syts. We validate our tools using both loss-of-function and gain-of-function approaches in live cells and further establish that our fluorescent lipids are bona fide E-Syt substrates using in vitro reconstitution in liposome transport assays. This strategy represents a purpose-driven tool for directly visualizing interorganelle lipid transport between the PM and the ER and establishes a framework to motivate development of additional analogous fluorescent probes for monitoring transport by shuttles and tubes at other MCS.

## RESULTS

Our approach for visualizing lipid transport at ER-PM contact sites is, essentially, a repurposing of bioorthogonal chemistry tools originally developed for visualizing phospholipase D (PLD) signaling activity. IMPACT (Imaging PLD Activity with Clickable Alcohols via Transphosphatidylation) involves metabolic labeling of cells with bioorthogonal primary alcohols that are used by PLDs in a transphosphatidylation reaction of phosphatidylcholine (PC) to form bioorthogonally labeled phosphatidyl alcohol lipids, followed by click chemistry tagging to generate fluorescent phosphatidyl alcohols^31–35^ (**Figure 1A**). By controlling the PLD stimulus during the metabolic labeling step and using rapid and fluorogenic inverse electron-demand Diels-Alder (IEDDA) *trans*-cyclooctene–tetrazine click chemistry tagging, the initial sites of fluorescent lipids can be restricted to desired membranes. For example, stimulation with phorbol esters results in PM-localized PLD activity, and thus the fluorescent lipid reporters of PLD activity are generated in this membrane^34^.

**Figure 1.**
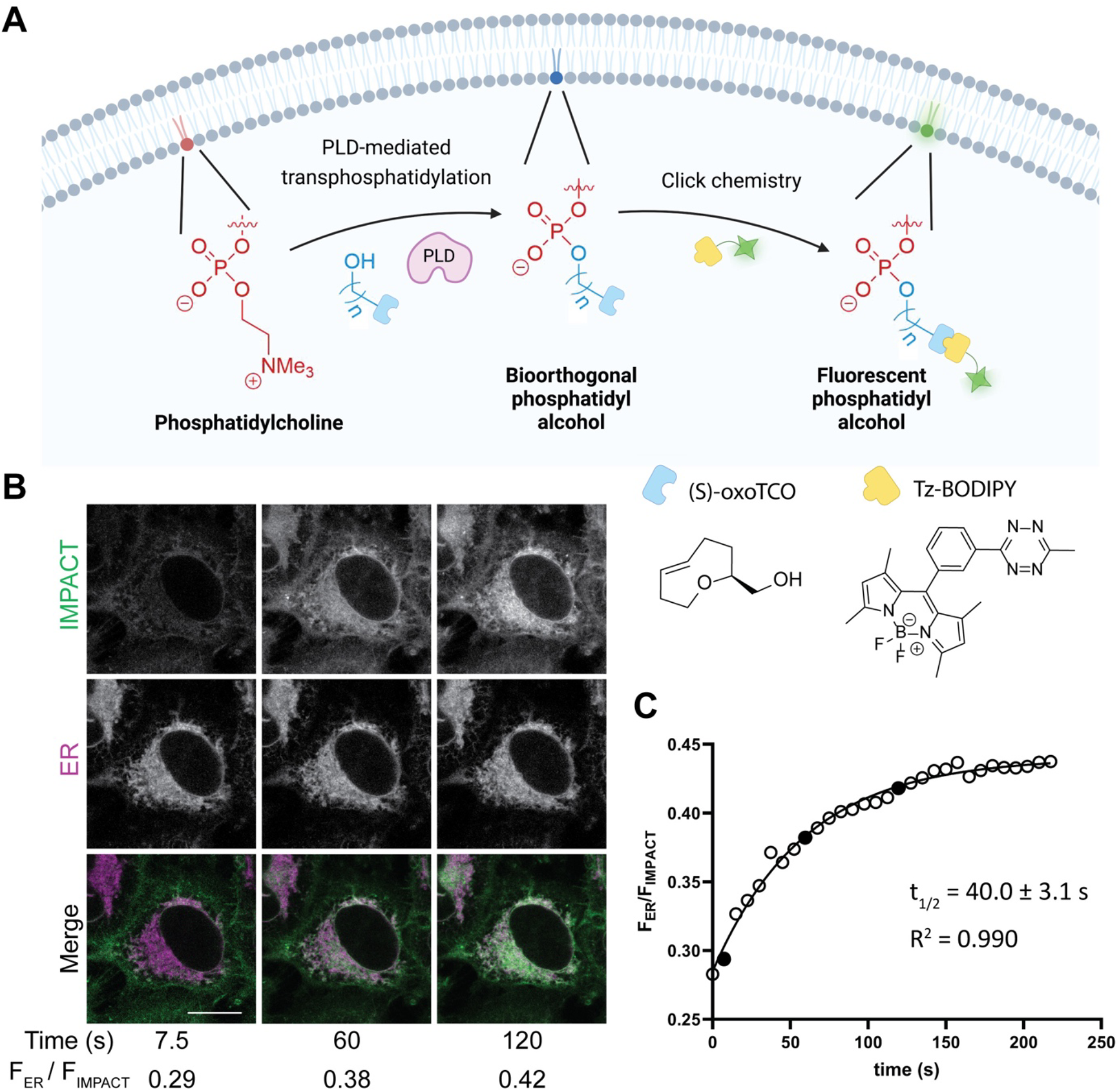
Fluorescent lipids generated by IMPACT are rapidly transported from the plasma membrane to the endoplasmic reticulum. (A) Schematic of Imaging Phospholipase D Activity with Clickable alcohols via Transphosphatidylation (IMPACT), a two-step strategy for visualizing phospholipase D (PLD) activity. Bioorthogonally tagged primary alcohols (e.g., with a *trans*-cyclooctene reagent (oxoTCO)) are first used as PLD transphosphatidylation substrates, and the subsequent phosphatidyl alcohols are fluorescently tagged using a bioorthogonal reaction (e.g., the inverse electron-demand Diels-Alder reaction with a fluorogenic tetrazine-BODIPY reagent (Tz-BODIPY)) to generate fluorescent lipid reporters of PLD activity. (B) IMPACT-derived fluorescent lipids generated at the PM by phorbol ester stimulation are rapidly transported to the ER. HeLa cells were treated with ER-Tracker Red, and transphosphatidylation was carried out with oxoTCO in the presence of phorbol 12-myristate-13-acetate (PMA) at 37 °C for 5 min, followed by a brief rinse and time-lapse imaging following Tz-BODIPY addition. Shown are representative images at indicated timepoints after addition of Tz-BODIPY (see also Video S1). IMPACT is shown in green, ER marker is shown in magenta, and colocalization appears as white. Ratios of IMPACT-derived fluorescence that overlaps with ER (F_ER_) relative to total IMPACT fluorescence (F_IMPACT_) were calculated for each timepoint. (C) Exponential fitting of kinetics of PM-to-ER trafficking. F_ER_/F_IMPACT_ was plotted versus time. Circles are raw data points, and the curve indicates an exponential fit. Filled circles correspond to F_ER_/F_IMPACT_ ratios for the timepoints shown in (B). R^2^, coefficient of determination of the fit; t_1/2_, average half-life of internalization (n=16 cells). Scale bar: 15 μm.

During the development of these “real-time” IMPACT tools, we noted the rapid translocation of fluorescent lipids on the second-to-minute timescale from the PM to the ER, prompting us to ask whether they might be reporting on LTP-mediated lipid transport at ER-PM contact sites. We began by performing real-time IMPACT labeling in HeLa cells by treatment with the oxo-*trans*-cyclooctene primary alcohol probe (oxoTCO) in the presence of phorbol 12-myristate 13-acetate (PMA) to stimulate endogenous PLD activity^36^, resulting in formation of oxoTCO-containing phosphatidyl alcohol lipids via PLD-mediated transphosphatidylation. After a brief rinse, the oxoTCO lipids were tagged via the IEDDA click reaction with a fluorogenic tetrazine–BODIPY reagent (Tz-BODIPY), resulting in formation of fluorescent phosphatidyl alcohol lipids. Consistent with our previous results, these fluorescent lipids appeared initially at the PM and then, over a few minutes, translocated to the ER (**Figure 1B**).

To quantify the PM-to-ER transport of the IMPACT-derived fluorescent lipids, we performed time-lapse imaging of RT-IMPACT fluorescence in cells co-stained with ER-Tracker Red. A plot of Pearson’s correlation coefficient between the two fluorescent channels, showed increasing colocalization of RT-IMPACT and ER-Tracker over time, enabling calculation of a half-time for PM-to-ER transport (**Figure 1C** and **Video S1**).

To assess a potential role for the E-Syts in this lipid trafficking event, we performed siRNA-mediated E-Syt1/2/3 triple knockdown (TKD) of the three human E-Syts (**Figure S1A**) and found that, relative to negative control siRNA, E-Syt TKD led to a statistically significant increase in transport half-time (14.4 ± 1.2 s for control siRNA vs. 21.5 ± 1.4 s for E-Syt TKD) (**Figure 2A–B** and **Videos S2–3**). To determine if the effects of the E-Syt siRNA were on-target, we next performed rescue experiments where E-Syt TKD-treated cells were transfected with an miRFP-tagged, siRNA-resistant form of E-Syt1. We found that miRFP-E-Syt1 expression fully reversed the effect of E-Syt TKD (restoring transport half-lives to ~15 s), indicating that the effect on IMPACT lipid transport rate induced by E-Syt TKD was in fact due to loss of E-Syt proteins (**Figure 2C–D and Video S4**).

**Figure S1.**
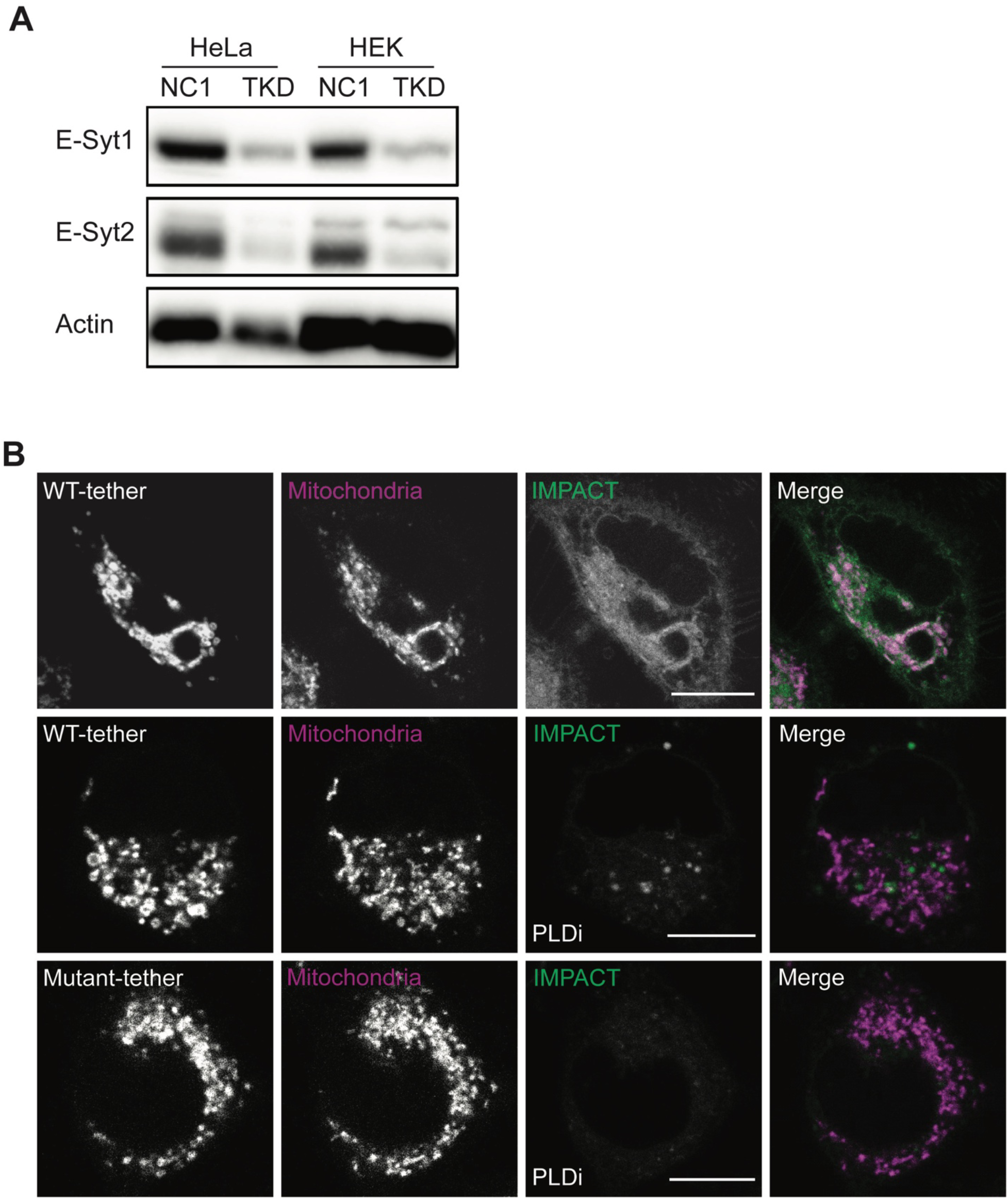
Validation of E-Syt RNAi and control experiments for IMPACT studies from Figure 3C. (A) Western blot analysis showing efficiency of E-Syt1/2/3 RNAi on the expression levels of E-Syt1 and E-Syt2 in HeLa and HEK 293 cells. Cells were transfected with siRNAs or control and harvested after 72 h, followed by Western blot analysis of lysates using anti-E-Syt1 and E-Syt2 antibodies. (B) Confocal microscopy analysis showing specificity of IMPACT to PLD activity in cells expressing WT-tether and Mutant-tether E-Syt constructs. HeLa cells were transfected with WT-tether (top) or Mutant-tether (bottom). After 24 h cells were treated with DMSO vehicle (top row) or the PLD1/2 inhibitor FIPI^37^ in DMSO (PLDi, bottom two rows) for 30 min, followed by incubation with a combination of PMA and oxoTCO to initiate IMPACT labeling, HaloTag-JF635 to label the E-Syt tether protein, and MitoView 405 to label mitochondria, at 37 °C for 5 min. Cells were rinsed for 1 min and labeled with Tz-BODIPY to complete IMPACT labeling for 1 min, followed by imaging by confocal microscopy. Note that addition of PLDi prevents substantial BODIPY fluorescence, indicating the specificity of the IMPACT labeling reagents to endogenous PLD activity. For clarity, the merged images indicate overlay of IMPACT (green) and MitoView 405 (magenta) channels, and colocalization in merged images appear as white. Scale bars: 10 μm.

**Figure 2.**
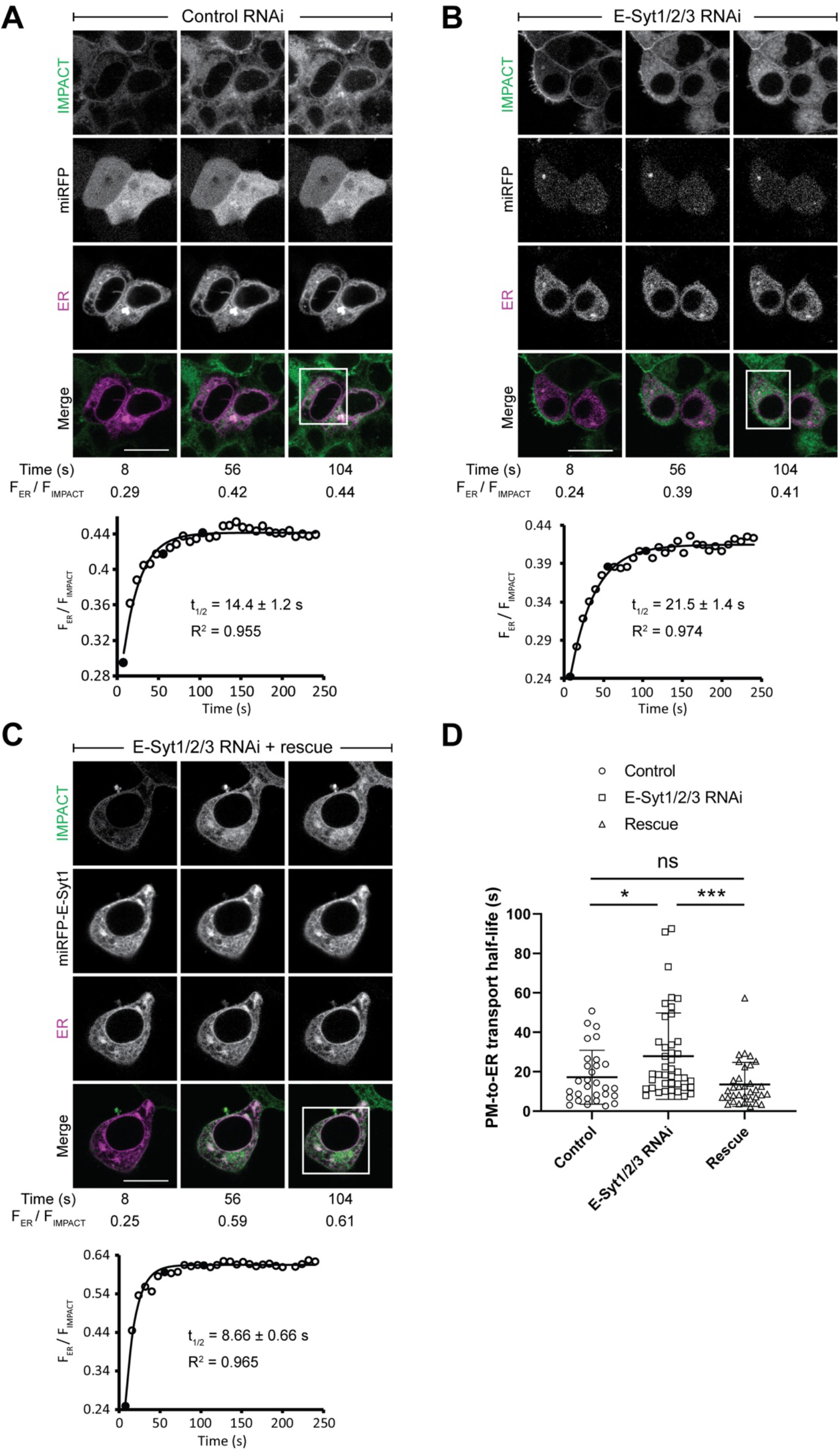
PM-to-ER trafficking of IMPACT-derived lipids depend upon the extended synaptotagmins (E-Syts). (A–C) Representative snapshots from time-lapse confocal imaging to assess the PM-to-ER trafficking kinetics of IMPACT-derived lipids (see also Videos S2–4). HEK 293 cells were first treated with control RNAi (A) or E-Syt1/2/3 RNAi (B–C), and after 48 h, cells were transiently transfected with Sec61β-mRFP (ER marker) and either miRFP empty vector (A–B) or miRFP-E-Syt1 (C). After 24 h, cells were labeled by IMPACT by treatment with PMA and oxoTCO for 5 min, rinse, and then time-lapse imaging in the presence of Tz-BODIPY. Shown in A–C are representative images taken 8, 56, and 104 s after addition of Tz-BODIPY, and merged images show only IMPACT (green) and ER marker (magenta) for clarity. Ratios of IMPACT-derived fluorescence that overlaps with ER (F_ER_) normalized to total IMPACT fluorescence (F_IMPACT_) were calculated for each time point and plotted below each set of images, with circles showing raw data points and curves showing exponential fits. Filled circles correspond to the three F_ER_/F_IMPACT_ values for the images shown above. R^2^ indicates the coefficient of determination of the fit and t1/2 indicates average half-life of internalization. Scale bars: 10 μm. (D) Scatter plot showing quantification of PM-to-ER transport half-lives of IMPACT-derived lipids. Each data point represents a single transfected cell (n=15–30 total cells from 3 dishes on different days). Mean and standard deviation are shown. One-way ANOVA, Games-Howell *post hoc* test: *, p < 0.05; **, p < 0.01; ***, p < 0.001; ns, not significant.

Because E-Syts are both LTPs and tethers functioning at ER-PM contact sites, it may be difficult to attribute the effects of endogenous E-Syt expression on the observed trafficking rates to their lipid transfer activities or their tethering functions, which promote the formation of contact sites. We sought to further dissect these two possibilities by reconstituting ectopic E-Syt-mediated lipid transfer in cells between the ER and a membrane other than the PM. Because we observed minimal localization of RT-IMPACT-derived lipids to mitochondria, we engineered an E-Syt-based artificial tether at ER-mitochondria contact sites to examine whether RT-IMPACT-derived lipids could be redirected from the ER (where they would accumulate due to endogenous E-Syt activity) to the mitochondria.

Our artificial E-Syt contained an N-terminal HaloTag and retained several N-terminal domains from E-Syt1, including the ER-targeting hydrophobic motif, the key SMP domain for mediating lipid transfer, and the C2A and C2B domains that mediate general phospholipid binding^38^. However, we replaced its C-terminal C2C, C2D, and C2E domains, which collectively enable PM recognition via Ca^2+^-dependent PI(4,5)P2 binding^39^, with an outer mitochondrial membrane targeting transmembrane helix sequence from OMP25, and we named this Halo-E-Syt1-mito construct as “WT-tether” (**Figure 3A**). To separate tethering function from LTP activity, we generated a mutant form (Halo-E-Syt1*-mito, termed “Mutant-tether”, bearing two point mutations in the SMP domain (V169W and L308W) that eliminate lipid binding (and therefore lipid transfer activity)^29^ (**Figure 3A**). Expression of WT-tether and Mutant-tether in HeLa cells revealed colocalization of both artificial tethers with both ER and mitochondrial membranes and minimal apparent perturbation to organelle morphology, indicating that these tethers localize to ER-mitochondria contact sites (**Figure 3B**).

**Figure 3.**
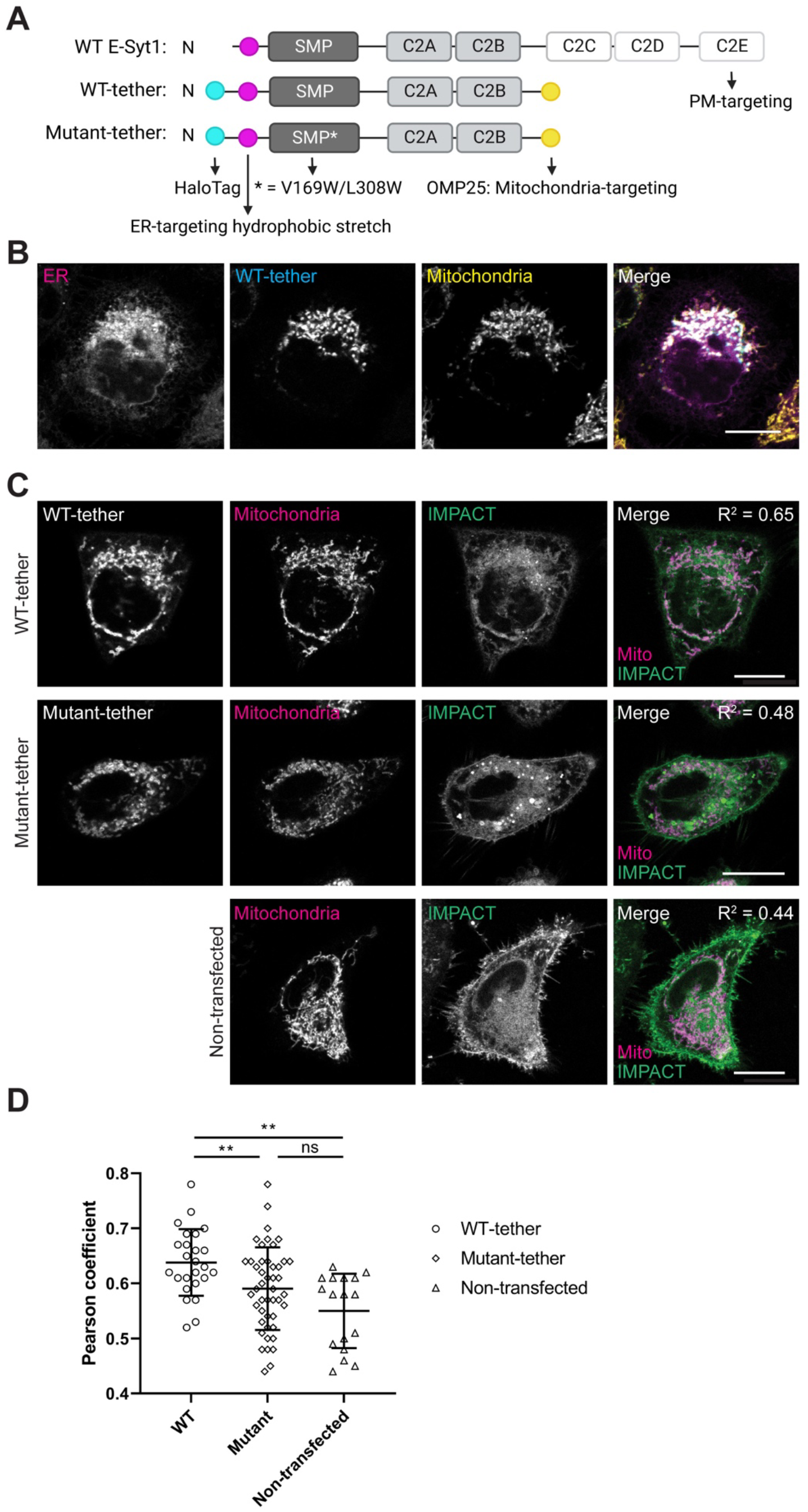
An artificial E-Syt1 tethered to ER-mitochondria contact sites redirects IMPACT-derived lipid transport toward mitochondria. (A) Domain maps of WT E-Syt1 and engineered constructs wherein a functional (WT-tether) or mutant (Mutant-tether) E-Syt1 lipid transfer module (SMP domain) is tethered at ER-mitochondria contact sites. (B) Confocal microscopy images of HeLa cells transfected with WT-tether and then stained with ER-Tracker Green (ER) and MitoView 405 (Mitochondria), showing colocalization of WT-tether with both organelles. Merged image shows overlay of all three channels: ER (magenta), WT-tether (blue), and mitochondria (yellow), with colocalization appearing as white. (C) Confocal microscopy analysis showing colocalization of RT-IMPACT-derived lipids with mitochondria. HeLa cells were transfected with WT-tether (top row), Mutant-tether (middle row), or no transfection (bottom row). After 24 h, cells were labeled via IMPACT by treatment with PMA and oxoTCO in the presence of HaloTag-JF635 and MitoView 405 to label WT-tether and stain mitochondria, respectively, for 5 min. After a brief rinse, cells were labeled with Tz-BODIPY for 1 min, followed by immediate imaging. Merged images show overlay of IMPACT (green) and MitoView 405 (magenta), with colocalization appearing as white. Pearson correlation coefficients (R^2^) were calculated for the two channels in the merged images. (D) Quantification of effects of WT-tether and Mutant-tether expression on trafficking of IMPACT-derived fluorescent lipids to mitochondria. Scatter plot shows quantification of Pearson coefficients from (C). For the WT-tether and Mutant-tether groups, n=25–46 individual transfected cells from three dishes acquired on separate days. For the Nontransfected group, each data point represents the average fluorescence of 15–30 cells from a single dish (n=17 dishes). Shown are mean and standard deviation. One-way ANOVA, Games-Howell post hoc test: *, p < 0.05; **, p < 0.01; ***, p < 0.001; ns, not significant. Scale bars: 10 μm.

To examine the ability of the artificial E-Syt constructs to mediate ER-to-mitochondrial trafficking of fluorescent lipids, we performed RT-IMPACT labeling in cells expressing either WT-tether or Mutant-tether and co-stained with markers of both the ER (ER-Tracker Red) and mitochondria (MitoView 405). We found that expression of WT-tether led to an increase in colocalization of RT-IMPACT and MitoView fluorescence relative to both untransfected cells and cells transfected with Mutant-tether (**Figures 3C** and **S1B**).

These studies indicate that artificially tethering the ER and mitochondria with an E-Syt-like molecule containing a functional SMP domain facilitates the transport of RT-IMPACT-derived fluorescent phosphatidyl alcohols from the ER to the mitochondria (after their rapid transport from PM to ER by endogenous E-Syts and other LTPs). Collectively, the cell-based loss-of-function and gain-of-function studies suggest that E-Syts can mediate the interorganelle lipid transfer of fluorescent lipids generated by IMPACT.

To directly assess whether E-Syts can transport IMPACT-derived fluorescent lipids between membranes, we reconstituted E-Syt-mediated LTP activity in vitro. For these studies, we expressed and purified E-Syt1 constructs wherein the N-terminal hydrophobic ER anchor was replaced by a 6xHis tag to enable both purification by metal affinity chromatography and anchoring onto liposomes containing Ni^2+^-chelating lipids. Two forms of E-Syt1 were generated: one containing the wildtype SMP domain and one containing the lipid-binding deficient V169W/L308W double mutant SMP domain.

In parallel, we chemoenzymatically synthesized a BODIPY-tagged dioleoyl phosphatidyl alcohol lipid by performing PLD transphosphatidylation on dioleoylphosphatidylcholine with a bioorthogonal alcohol followed by click chemistry tagging to append BODIPY and purification by HPLC^34^. Finally, we prepared two populations of liposomes. The “donor” liposomes containing the Ni^2+^-chelating lipids, the IMPACT-derived BODIPY lipids, and rhodamine-labeled PE to quench the BODIPY lipids by FRET. The “acceptor” liposomes contained anionic plasma membrane lipids including phosphatidylserine and, crucially, PI(4,5)P2. Incubation of both liposome populations with the 6xHis-E-Syt1 constructs should lead to tethering of the liposomes and, in the case of the E-Syt1 construct with the wildtype SMP domain, lead to Ca^2+^-dependent phospholipid transfer between the liposomes, causing dilution of the fluorescent lipids, dequenching, and therefore an increase in BODIPY fluorescence as assessed by a fluorescence plate reader (**Figure 4A**).

**Figure 4.**
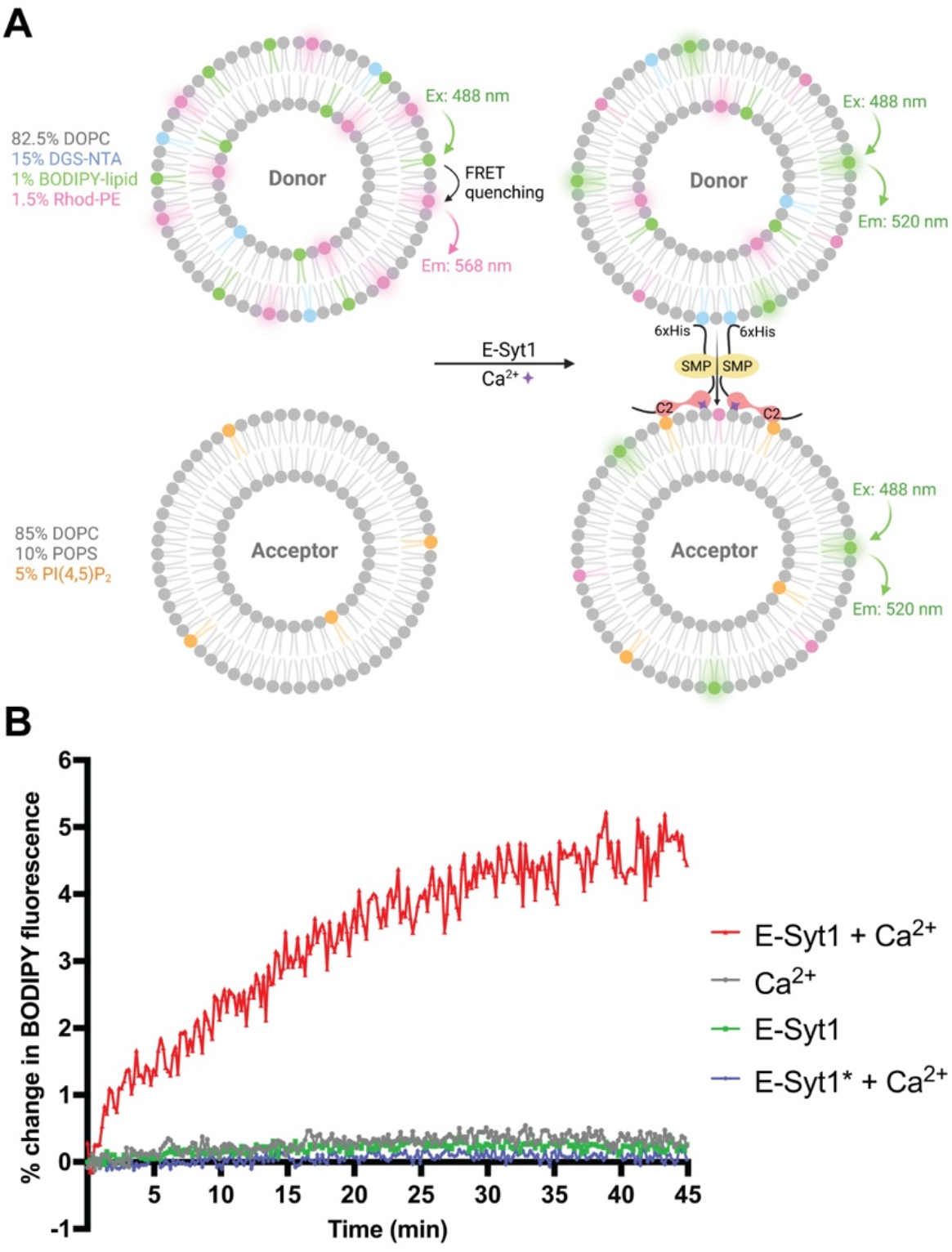
IMPACT-derived BODIPY lipids can be transported by E-Syt1 in vitro. (A) Schematic of the FRET-based in vitro lipid transfer assay. Donor and acceptor liposomes (compositions indicated in the figure) are tethered by E-Syt1, and an increase in BODIPY fluorescence, the result of FRET dequenching following E-Syt1 and Ca^2+^-dependent transport from donor to acceptor liposomes, is monitored using a microplate reader. (B) Time course of normalized BODIPY fluorescence signal from mixtures of liposomes incubated at room temperature with the indicated combination of E-Syt1, E-Syt1* (mutant deficient in lipid transfer), or no protein in the presence or absence of Ca^2+^. For all experiments, the donor:acceptor liposome ratio was 1:4 and the overall protein:lipid ratio was 1:1000. Shown are representative data from a single experiment (n=3 replicate experiments were performed that all yielded similar results).

Indeed, we observed the appearance of increasing BODIPY fluorescence over time when the WT 6xHis-E-Syt1 construct was incubated with donor and acceptor liposomes (**Figure 4B**). Importantly, no fluorescence increase was observed in three negative control samples that lacked Ca^2+^ or contained the E-Syt1 construct with the mutant SMP domain with or without Ca^2+^ addition. We conclude from these studies that IMPACT-derived fluorescent phosphatidyl alcohols, like NBD-functionalized lipids and several types of native glycerolipids, are bona fide lipid transfer substrates of E-Syt1. Collectively, the cellular imaging and in vitro reconstitution studies here indicate that IMPACT can serve as a live-cell readout of lipid transfer from the PM to the ER and that a fraction of this transport activity revealed by IMPACT is accomplished by the E-Syt proteins.

## DISCUSSION

Fluorescent probes are invaluable tools for visualizing organelle membranes, membrane trafficking events, and flux through lipid biosynthetic pathways in live cells. A key aspect of lipid biosynthesis is the often-rapid transport of lipids between different organelles, and non-vesicular pathways are now appreciated as the major contributors to intracellular lipid flux. Such non-vesicular transport is largely mediated by LTPs functioning at MCS as tethers and/or transporters. Structural studies and in vitro biochemical reconstitution of transport activity have revealed invaluable information about the substrate specificity, directionality, and energy requirements for individual LTPs. However, it remains unclear how such parameters determined in vitro may translate to native cellular settings, and the available approaches for assaying LTP function in cells rely on indirect readouts, offering much less molecular and quantitative information compared to in vitro approaches.

Here, we began with the observation that fluorescent lipids deriving ultimately from PLD activity at the PM were rapidly internalized to the ER. Supporting these observations were the knowledge that phorbol ester stimulation results in PLD activation at the PM and that a rapid, two-step metabolic labeling and bioorthogonal tagging approach, IMPACT, would enable generation of fluorescent phospholipid analogs at the PM prior to substantial redistribution of lipids. The accumulation of IMPACT-derived lipids at the ER as the first post-PM station, and the absence of any endosomal IMPACT fluorescence at these early timepoints, strongly suggested the involvement of non-vesicular lipid transport occurring via MCS.

Using a candidate-based approach, we identified the E-Syts as major players in this PM-to-ER transport of the IMPACT-derived fluorescent lipids. Extensive structural and biochemical studies on E-Syts have established that these proteins bind to glycerolipids via their hydrophobic regions within the SMP domain, with little to no recognition of the polar head group. In addition, such transport by E-Syts is bidirectional and ATP-independent, which renders the lipid transfer process thermodynamically driven. As a result, the lipid transfer activity of E-Syts not only serves as a means to remove specific metabolic lipid products (e.g., diacylglycerol) from the plasma membrane, but it may also have broader roles in maintaining lipid homeostasis for cells to cope with unnatural perturbations, such as doping PM with a large amount of unnatural fluorescent lipids derived from IMPACT labeling.

Using RNAi-mediated knockdown and rescue experiments, we demonstrated that cells react to IMPACT-derived unnatural fluorescent phospholipids by transporting them from PM to ER, in large part via E-Syts. The residual transport activity in the absence of E-Syts indicates that other LTPs at ER-PM contacts play a role, underscoring the complexities and redundancies built into this important system. Further, we engineered an artificial tether made by replacing the C-terminal PM-targeting domains of E-Syts with a mitochondria-targeting sequence. This tether localized to ER-mitochondria contacts and led to enhanced trafficking of unnatural lipids toward mitochondria, downstream of their transport from the PM to ER by endogenous E-Syts in these studies. Finally, we reconstituted the lipid transfer activities of E-Syts in vitro with purified proteins in liposome assays, demonstrating that IMPACT-derived fluorescent lipids are bona fide substrates of the E-Syt SMP domain.

In sum, these data support that E-Syts can transport unnatural fluorescent lipids that are rapidly generated on the cytosolic leaflet of the PM, using PLD transphosphatidylation and bioorthogonal tagging, to the ER. As such, IMPACT constitutes a tool for directly monitoring PM-to-ER interorganelle lipid transport by E-Syts and other LTPs. Such broad substrate specificity of E-Syts towards unnatural lipids with bulky head groups is intriguing and suggests that these LTPs may be capable of transporting a wide array of glycerolipids in cells depending on experimental perturbations, to restore lipid homeostasis. Beyond E-Syts and ER-PM contact sites, we envision that our approach is generalizable to LTPs with broad substrate scope — i.e., those that are head group-agnostic — acting at other membrane contact sites. Recent work from our lab has established optogenetic PLDs whose localization and activity are controllable by light and are therefore not restricted to locations visited by endogenous PLDs. Thus, we envision that this work will pave the way for a collection of IMPACT-based and -inspired tools for the precise installation of fluorescent phospholipid reporters on multiple target membranes, to aid in the search for new LTPs and characterization of their transport activities within live cells.

## ACKNOWLEDGMENTS

We acknowledge support from the NSF (CAREER CHE-1749919), a Beckman Young Investigator Award, and a Sloan Research Fellowship. We thank Reika Tei and Kane Wu for technical assistance, Pietro De Camilli and Gunther Hollopeter for sharing E-Syt1 and HaloTag plasmids respectively, and the Fromme lab for generously sharing reagents and equipment.

## AUTHOR CONTRIBUTIONS

D.L., L.L., and J.M.B. designed experiments, analyzed results, and wrote the manuscript. D.L. performed all cell-based experiments, and L.L. performed in vitro liposome experiments. D.L. and L.L. contributed equally.

## MATERIALS AND METHODS

### Reagents

E-Syt1 siRNA duplexes were purchased from Dharmacon (Cat # J-010652-06-0010). E-Syt2 and E-Syt3 siRNA duplexes were purchased from Integrated DNA Technologies (Cat # HSC.RNAI.N020728.12.5 and HSC.RNAI.N031913.12.6 respectively). Antibodies to E-Syt1, E-Syt2, and E-Syt3 were purchased from Sigma-Aldrich (Cat # HPA016858, HPA002132, and HPA039200, respectively). Lipofectamine 2000 (Cat # 11668019) and Lipofectamine RNAiMAX (Cat # 13778150) were purchased from Thermo Fisher. FIPI was purchased from Cayman Chemical (Cat # 13563). Phorbol 12-myristate 13-acetate (PMA) was purchased from ChemCruz (Cat # SC-3576). MitoView 405 was purchased from Biotium (Cat # 70070). ER-Tracker Green (Cat # E34251) and ER-Tracker Red (Cat # E34250) were purchased from Thermo Fisher. HaloTag-JF635 was synthesized according to previous protocols^40^. RT-IMPACT reagents, namely oxoTCO and Tz-BODIPY, were synthesized as previously reported^41,42^. See Table S1 for details.

### Cell culture

HEK 293 and HeLa cells were grown in DMEM (Dulbecco’s modified Eagle medium) supplemented with 10% FBS, 1% penicillin/streptomycin (HEK 293 cells were additionally supplemented with 1 mM sodium pyruvate) were maintained in a 5% CO2, moisture-saturated atmosphere at 37 °C. Cell densities were maintained between 10^5^ and 1.6 x 10^6^ cells/mL. For cell labeling experiments, all buffers or media were warmed to 37 °C or room temperature prior to addition to cells, and incubations were done at 37 °C.

### Primers and cloning

EGFP-E-Syt1 plasmid ^43^ was obtained from the De Camilli lab (Yale University). In all PCR reactions, Phusion High-Fidelity DNA Polymerase (New England Biolabs, Cat # M0530S) was used for amplifying fragments. PfuUltra High-fidelity DNA Polymerase (Agilent Technologies, Cat # 600382) was used for site-directed mutagenesis. Gibson assembly was performed using Gibson Assembly HiFi 1 Step Starter Kit (Synthetic Genomics, Cat # GA1100-S).

miRFP-E-Syt1 was cloned by PCR and ligated by a three-component Gibson assembly. The miRFP fragment was amplified from pmiRFP-N1 (Addgene Cat # 79987) by the primers Gib-miRFP-F (taagcttggtaccgagctcgATGGTAGCAGGTCATGCCTC) and Gib-miRFP-AgeI-(E-Syt1)-R (atcgctccataccggtGCTCTCAAGCGCGGTGATC). The E-Syt1 fragment was amplified from the EGFP-E-Syt1 plasmid by Gib-AgeI-E-Syt1-(miRFP)-F (ccgcgcttgagagcaccggtATGGAGCGATCTCCAGGAG) and Gib-E-Syt1-R (tccaccacactggactagtgCTAGGAGCTGCCCTTGTC). pcDNA5-FRT-TO (ThermoFisher Cat # V652020) was digested by BamHI. The digested plasmid was then ligated with miRFP and E-Syt1 fragments by Gibson assembly.

An RNAi-resistant form of miRFP-E-Syt1 produced by introducing six silent point mutations in the region targeted by the RNAi oligos using site-directed mutagenesis with the following primers (targeted nucleotides are shown in uppercase: E-Syt1_siRNA-res_S, ctgggccaggtgaaactgactctgtggtaTtaTTCCgaGgagcgaaagctggtcagcattgttc; E-Syt1_siRNA-res_AS, gaacaatgctgaccagctttcgctcCtcGGAAtaAtaccacagagtcagtttcacctggcccag)^43^.

Halo-E-Syt1 was cloned by PCR and ligated by a two-component Gibson assembly. The Halo-TEV fragment was amplified from HaloTag-TEVsite_pcDNA5.FRT Plasmid (obtained from Hollopeter lab, Cornell University)^44^ by the primers (AgeI)-GFP-C1-AgeI-Halo-F (cgtcagatccgctagcgctaaccggtgccaccatggcagaaatcggtactg) and (SalI)-GFPC1-GGSG-Halo-R (ctcctggagatcgctccatgcctgatccaccgttatcgctctgaaagtacag). The EGFP-E-Syt1 plasmid was digested using AgeI and SalI. The digested plasmid was then ligated together with the Halo-TEV fragment using Gibson assembly.

Halo-E-Syt1-OMP25 was cloned by PCR and ligated by a two-component Gibson assembly. The GGS-OMP25-MCS fragment was amplified from thee mCherry-OMP25 plasmid (obtained from De Camilli lab, Yale University) using the primers OMP25-MCS_fwd (gccccacctcgacccagatctcgagctcaaggtg) and OMP25-MCS_rev (tttgtgaaatttgtgatgctattgctttatttgtaaccattataagc). The AgeI-E-Syt1-C2AB fragment was amplified from the Halo-E-Syt1 plasmid using AgeI-E1-C2AB_F (gcatcacaaatttcacaaataaagcatttttttc) and AgeI-E1-C2AB_R (gggtcgaggtggggcatc). These two fragments were then ligated together using Gibson assembly.

Halo-E-Syt1*-OMP25 was further cloned by site-directed mutagenesis using primer sets E-Syt1_L308W_MF (tgccttcctcgtgttgcccaaccgatggctggtgccccttgtgcctgaccttcaagatgt), E-Syt1_ L308W _MR (acatcttgaaggtcaggcacaaggggcaccagccatcggttgggcaacacgaggaaggca), E-Syt1_V169W_MF (tctggctgaaactgtggctccggcttggaggggatctaacccccatctgcaaacatttac) and E-Syt1_V 169W_MR (gtaaatgtttgcagatgggggttagatcccctccaagccggagccacagtttcagccaga)^29^.

### General imaging methods

All imaging experiments were performed on a Zeiss LSM 800 confocal laser scanning microscope equipped with a 40X 1.4 NA Plan Apochromat objective, 405, 488, 561, and 640 nm solid-state lasers, and two GaAsP PMT detectors, using the Zeiss Zen Blue 2.3 software. All image analysis was performed using Fiji/ImageJ.

### Fluorescence colocalization imaging of IMPACT with organelle markers

HeLa or HEK cells (150,000 cells) were seeded on 35-mm glass-bottom imaging dishes (MatTek or Matsunami) for 24 h prior to experiments. Where indicated, the cells were then transfected with RNAi using Lipofectamine RNAiMAX (Thermo Fisher, 24 h after seeding), and/or the indicated plasmid with Lipofectamine 2000 (Thermo Fisher, 24 h after RNAi treatment or 24 h before imaging/harvesting). 72 h after RNAi treatment (or 24 h after plasmid transfection, if any), cells were first treated with the PLD inhibitor FIPI (PLDi, 750 nM)^37^, or DMSO vehicle in media at 37 °C. Cells for Western blot analysis were also harvested after 72 h of RNAi treatment. Subsequently, for imaging experiments, freshly prepared oxoTCO (3 mM) together with PMA (100 nM), organelle markers (ER-Tracker Red and MitoView 405), and PLD inhibitor/DMSO in media (100 μL) were carefully added to cover the central glass well.

(*Cautionary note*: oxoTCOs are reported to have limited water stability^41^. Therefore, all aqueous oxoTCO solutions (e.g., those in DMEM-containing media) were used within 20 min of their generation. For example, we dissolved oxoTCO in 200 μL of DMEM with PLD inhibitor/DMSO and respective stimulus and used it only for two consecutive replicates rather than making a single stock solution for an entire day of experiments at the beginning of the day.)

The dish was incubated for 5 min, the treatment media was aspirated, the cells were rinsed with PBS (1 mL) briefly, and the cells were then rinsed in DMEM (500 μL) for 1 min at 37 °C. Rinses after oxoTCO incubation were performed in DMEM supplemented with 10% FBS and 1% P/S (media). The media was aspirated and replaced with Tz-BODIPY (100 μL, 1 μM in PBS) for 1 min. Tz-BODIPY was then aspirated and replaced with Tyrode’s-HEPES buffer. Cells were imaged immediately afterwards. Multicolor images were obtained in two- or three-channel, line-switching mode (**Figures 3** and **S1**). Z stacks were taken with 0.45 μm sectioning.

### Real-time imaging of trafficking half-lives of IMPACT-derived fluorescent lipids

IMPACT using oxoTCO and Tz-BODIPY and was performed as described previously^34,35,45^. Freshly prepared oxoTCO (3 mM) together with PMA (100 nM) in media (100 μL) were carefully added to cover the central glass well of the imaging dish. The dish was incubated for 5 min, the treatment media was aspirated, the cells were rinsed with PBS (1 mL) briefly, and the cells were then rinsed in media (500 μL) for 1 min at 37 °C. The media was replaced with 100 μL of PBS buffer to cover the center of the glass bottom and the dish was mounted on the microscope. The cells to be imaged were quickly located, and time-lapse imaging with an interval of 8 s and duration of 4 min begun (as in **Figures 1–2**). Tz-BODIPY (100 μL, 1 μM in PBS) was added dropwise but quickly to the center of the dish during acquisition. The percentage of total IMPACT fluorescence (488 channel) colocalizing with a mask of ER-Tracker Red (561 channel) (F_ER_/F_IMPACT_) was determined in ImageJ for each time point. For **Figures 1C** and **2**, F_ER_/F_IMPACT_ vs. time was then fit to a mono-exponential decay function [y = A_1_ * exp(-x/t_1_) + y_0_] using Origin Pro 8, with the half-life equal to [ln(2) * t_1_]. For **Figure 3**, Pearson correlation coefficients between IMPACT and either ER marker or mitochondrial marker channels were determined in ImageJ. All half-lives and Pearson coefficients were plotted as scatter plots (**Figures 2D and 3D**) using GraphPad Prism 8.

### Synthesis of IMPACT-derived BODIPY-lipid for liposome assays

The BODIPY-lipid for FRET-based lipid transfer assay was synthesized by an in vitro PLD transphosphatidylation reaction followed by click chemistry tagging reaction as described previously^31,32^ but on a larger scale. PLD (1 mg of superPLD, clone 2-48^46^) and 500 mM azidopropanol were added to a solution of PBS (100 μL, pH 7.4) containing dioleoylphosphatidylcholine (DOPC, 80 μL of an 10-mg/mL solution in ethyl acetate). This transphosphatidylation reaction was allowed to proceed at 37 °C for 8 h, and the azide-containing phosphatidyl alcohol lipid product was isolated by a Bligh-Dyer extraction. To fluorescently tag this azido lipid, 1.6 mg of the crude residue from the combined organic fractions from the Bligh-Dyer extraction was dissolved in chloroform:methanol:water (73:23:3), and 2 μmol of BCN-BODIPY^47^ was added and the mixture was incubated for 20 h at 42 °C. The crude reaction product was analyzed and purified by HPLC using 3 μm, 250 x 4.6 mm (analytical, Phenomenex Luna) and 5 μm, 250 × 9.4 mm (semi-prep, Agilent) normal-phase silica columns, with a binary gradient elution system wherein solvent A was chloroform:methanol:ammonium hydroxide (95:7:0.5) and solvent B was chloroform:methanol:water:ammonium hydroxide (60:34:5:0.5). Detection wavelengths were 495 nm (excitation) and 510 nm (emission).

### Protein purification

The E-Syt1 proteins used for lipid transfer assays were purified as described previously^29^. Briefly, WT and V169W/L308W double mutant human E-Syt1 (93-1104 aa) with N-terminal 6xHis-tags (plasmid obtained from De Camilli lab at Yale University) were expressed in Expi293 suspension HEK cells. Cells were harvested 72 h post-transfection by centrifugation at 500 g for 10 min at room temperature, and the pellet was resuspended in lysis buffer (25 mM Tris-HCl pH 8.0, 300 mM NaCl, 0.5 mM TCEP, 10 mM imidazole, cOmplete EDTA-free protease inhibitor). Cells were lysed by 3 cycles of freezing with liquid nitrogen and thawing in a 37 °C water bath and clarified by centrifugation at 17,000 g for 30 min at 4 °C. Clarified lysate was incubated with TALON Metal Affinity Resin overnight, and resin was rinsed with wash buffer (25 mM Tris-HCl pH 8.0, 300 mM NaCl, 0.5 mM TCEP, 20 mM imidazole). The protein was eluted with elution buffer (25 mM Tris-HCl pH 8.0, 300 mM NaCl, 0.5 mM TCEP, 400 mM imidazole) and further concentrated using a 30 kDa molecular weight cutoff centrifugal concentrator.

### Liposome preparation

Lipids in chloroform were mixed to create acceptor and donor liposomes separately in the following ratios: 85% DOPC, 10% POPS, 5% PI(4,5)P2 for acceptor liposomes; 82.5% DOPC, 15% DGS-NTA, 1.5% BODIPY-lipid, 1% Rhod-PE for donor liposomes. Lipid mixtures were dried to thin films under a stream of N2 gas, placed under vacuum for 2 h, and resuspended in buffer containing 25 mM Tris-HCl, pH 8.0, 150 mM NaCl, 0.5 mM TCEP to a total lipid concentration of 2.5 mM. The suspension was subjected to 10 freeze-thaw cycles and extruded 11 times through a polycarbonate filter with 50-nm pore size in a mini-extruder.

### FRET-based lipid-transfer assays

Lipid transfer experiments were performed in black flat bottom 96-well plates, with 50 μL reaction volumes containing 0.5 μM protein (E-Syt1 or E-Syt1* as shown in **Figure 4A**) and 500 μM final total lipid concentration with a 1:4 ratio of donor:acceptor liposomes, either in the presence or absence of 100 μM CaCl2. Lipid transfer was initiated by adding protein and terminated by adding 10 μL of a 2.5% aqueous solution of n-Dodecyl-β-D-maltoside (DDM) to solubilize the liposomes. BODIPY fluorescence intensity was monitored with an excitation of 460 nm and emission of 538 nm every 11 s over 45 min at 25 °C using a BioTek Synergy H1 microplate reader. All data were baseline corrected by normalizing to the difference between the maximum (average of last 5 data points after adding DDM) and the minimum (average of first 3 data points after adding protein).

### Statistics and reproducibility

All imaging experiments show representative images from experiments performed in at least three biological replicates on different days, where each replicate refers to a dish of cells with approximately 15-30 cells in total and 2-10 transfected cells in the field of view. For imaging with transfected cells, measurements were made on individual transfected cells, with each transfected cell being a data point in **Figure 2D** (all samples) and **Figure 3D** (left two samples). For imaging experiments with non-transfected cells, each dish contained approximately 15-30 cells in total, and the parameters for all cells were averaged and recorded as one data point for **Figure 3D** (rightmost sample). Exact numbers of replicate experiments and sample sizes are provided in each figure legend. For **Figures 2D** and **3D**, p-values were calculated using one-way ANOVA, followed by Games-Howell post-hoc test (for samples of unequal variance).

## LEGENDS FOR SUPPLEMENTAL VIDEOS

**Video S1. Time-lapse imaging of PM-to-ER lipid transport in cells labeled via IMPACT (related to Figure 1).** HeLa cells were labeled as described in Figure 1B, and time-lapse imaging was performed by confocal microscopy. Images were acquired every 3 s, and Tz-BODIPY was added in between the first and second frames. Scale bar: 15 μm.

**Video S2 (related to Figure 2). Time-lapse imaging of cells treated with control RNAi showing PM-to-ER trafficking of IMPACT-derived lipids.** HEK 293 cells were labeled as described in Figure 2A, and time-lapse imaging was performed by confocal microscopy. Images were acquired every 3 s, and Tz-BODIPY was added in between the first and second frames. Scale bar: 10 μm.

**Video S3 (related to Figure 2). Time-lapse imaging of cells treated with E-Syt1/2/3 RNAi showing PM-to-ER trafficking of IMPACT-derived lipids.** HEK 293 cells were labeled as described in Figure 2B, and time-lapse imaging was performed by confocal microscopy. Images were acquired every 3 s, and Tz-BODIPY was added in between the first and second frames. Scale bar: 10 μm.

**Video S4 (related to Figure 2). Time-lapse imaging of cells treated with E-Syt1/2/3 RNAi and transfected with miRFP-E-Syt1 showing PM-to-ER trafficking of IMPACT-derived lipids.** HEK 293 cells were labeled as described in Figure 2C, and time-lapse imaging was performed by confocal microscopy. Images were acquired every 3 s, and Tz-BODIPY was added in between the first and second frames. Scale bar: 10 μm.

**TABLE S1.**
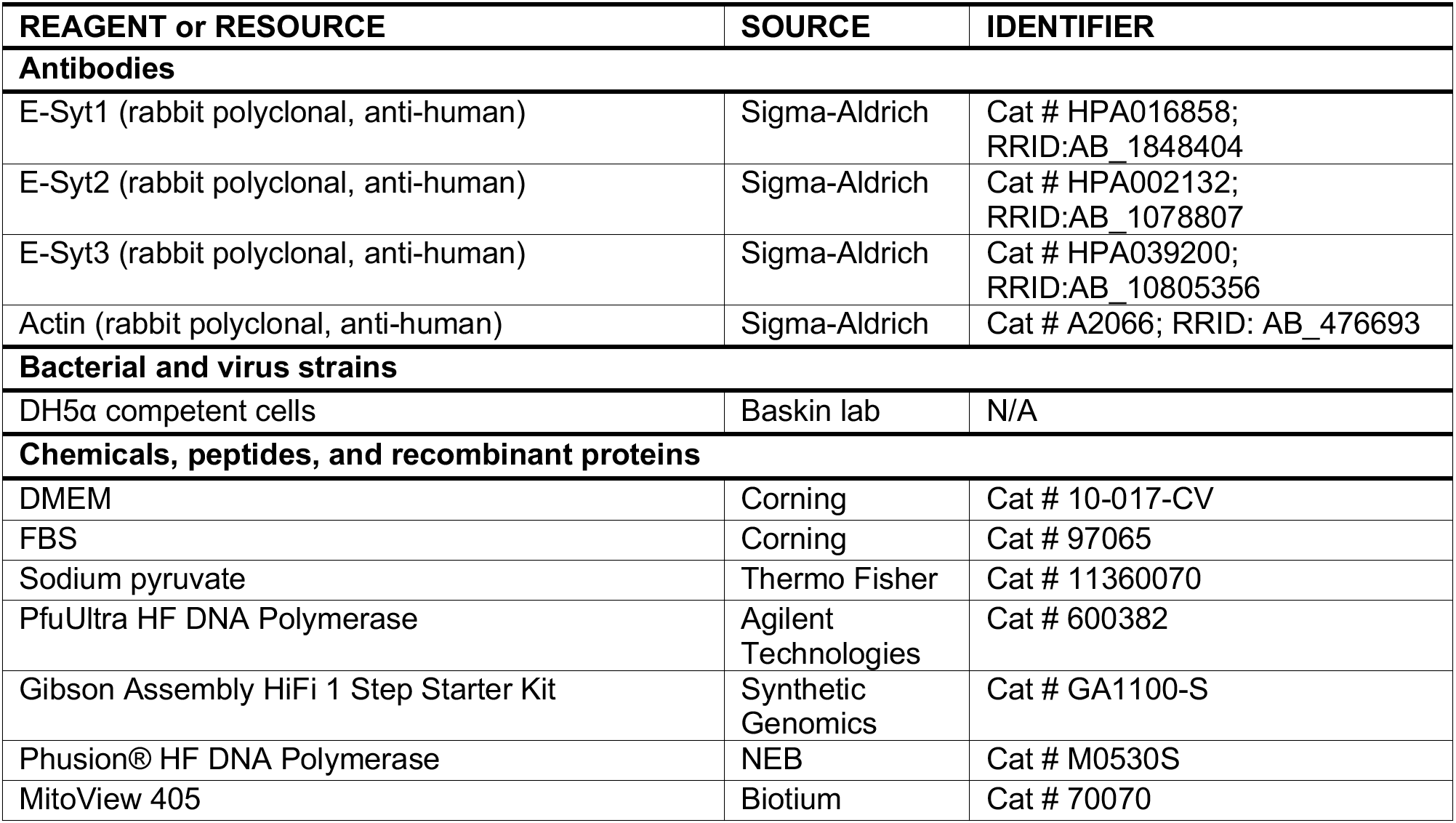
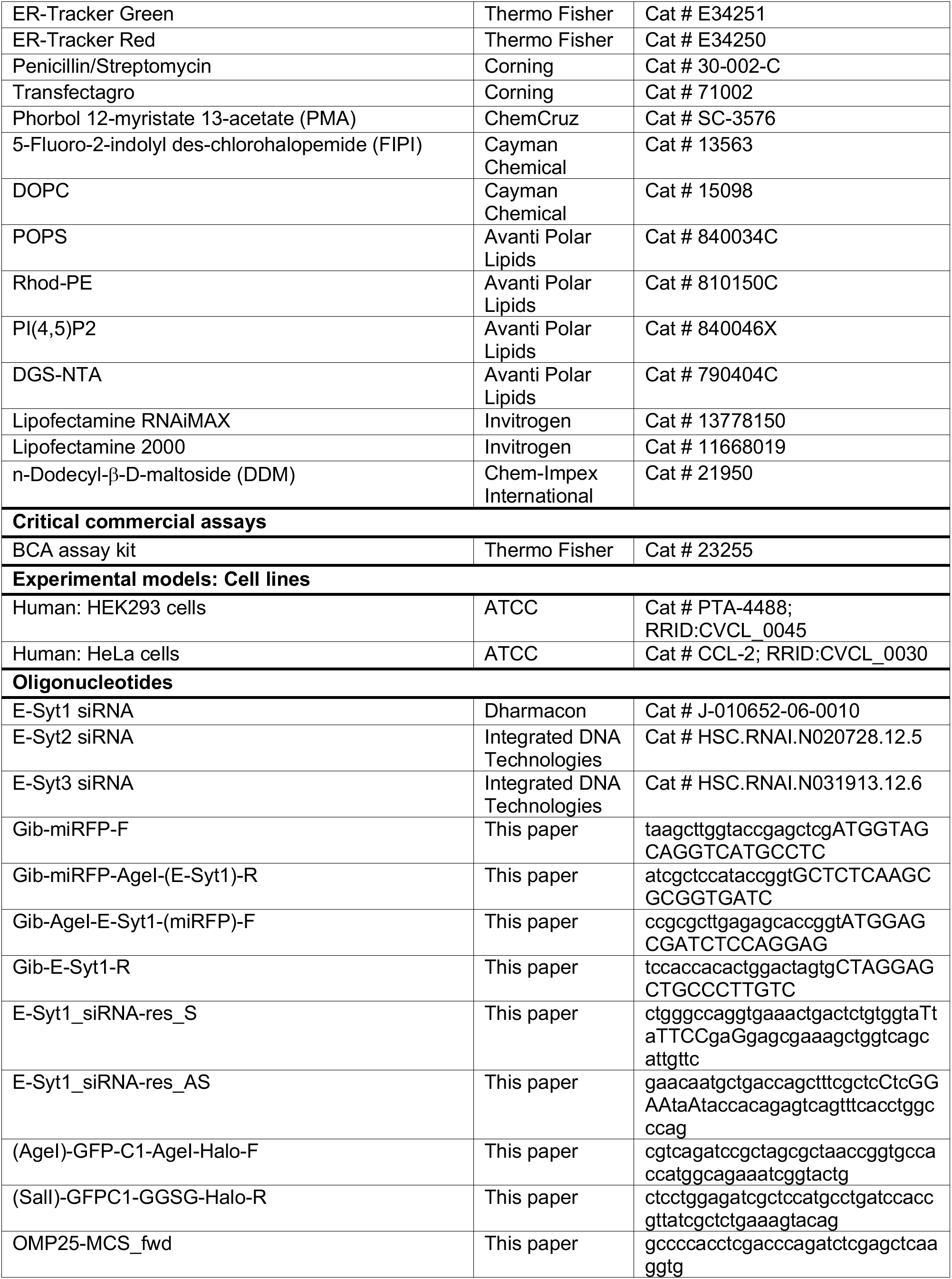
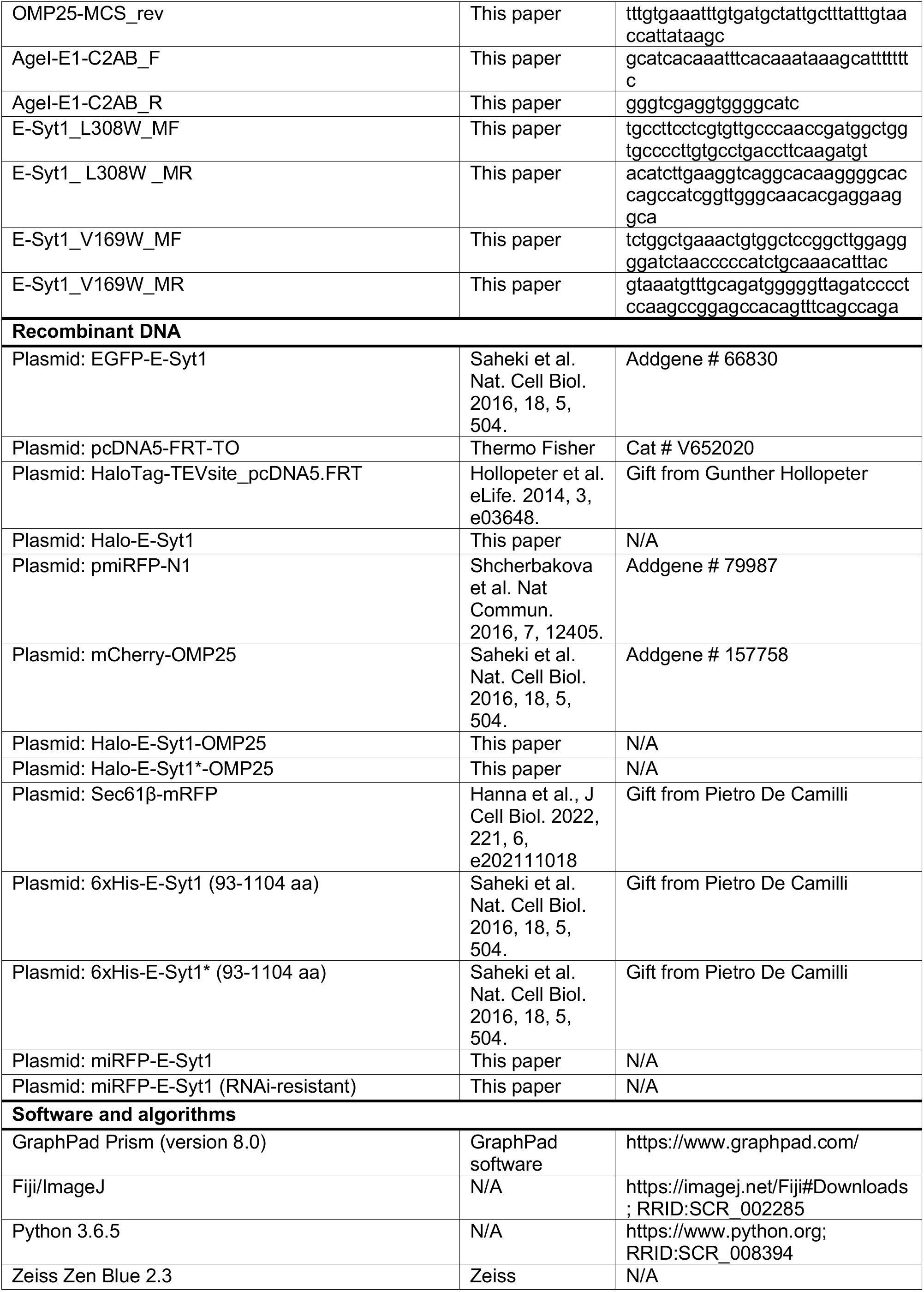
List of reagents used in this study.

## REFERENCES

1. Reinisch, K.M., and Prinz, W.A. (2021). Mechanisms of nonvesicular lipid transport. J. Cell Biol. 220, e202012058. 10.1083/jcb.202012058.

2. Wong, L.H., Gatta, A.T., and Levine, T.P. (2019). Lipid transfer proteins: the lipid commute via shuttles, bridges and tubes. Nat. Rev. Mol. Cell Biol. 20, 85–101. 10.1038/s41580-018-0071-5.

3. Kolehmainen, J., Black, G.C.M., Saarinen, A., Chandler, K., Clayton-Smith, J., Träskelin, A.-L., Perveen, R., Kivitie-Kallio, S., Norio, R., Warburg, M., et al. (2003). Cohen Syndrome Is Caused by Mutations in a Novel Gene, COH1, Encoding a Transmembrane Protein with a Presumed Role in Vesicle-Mediated Sorting and Intracellular Protein Transport. Am. J. Hum. Genet. 72, 1359–1369. 10.1086/375454.

4. Ionita-Laza, I., Capanu, M., Rubeis, S.D., McCallum, K., and Buxbaum, J.D. (2014). Identification of Rare Causal Variants in Sequence-Based Studies: Methods and Applications to VPS13B, a Gene Involved in Cohen Syndrome and Autism. PLOS Genet. 10, e1004729. 10.1371/journal.pgen.1004729.

5. Lesage, S., Drouet, V., Majounie, E., Deramecourt, V., Jacoupy, M., Nicolas, A., Cormier-Dequaire, F., Hassoun, S.M., Pujol, C., Ciura, S., et al. (2016). Loss of VPS13C Function in Autosomal-Recessive Parkinsonism Causes Mitochondrial Dysfunction and Increases PINK1/Parkin-Dependent Mitophagy. Am. J. Hum. Genet. 98, 500–513. 10.1016/j.ajhg.2016.01.014.

6. Gauthier, J., Meijer, I.A., Lessel, D., Mencacci, N.E., Krainc, D., Hempel, M., Tsiakas, K., Prokisch, H., Rossignol, E., Helm, M.H., et al. (2018). Recessive mutations in VPS13D cause childhood onset movement disorders. Ann. Neurol. 83, 1089–1095. 10.1002/ana.25204.

7. Seong, E., Insolera, R., Dulovic, M., Kamsteeg, E.-J., Trinh, J., Brüggemann, N., Sandford, E., Li, S., Ozel, A.B., Li, J.Z., et al. (2018). Mutations in VPS13D lead to a new recessive ataxia with spasticity and mitochondrial defects. Ann. Neurol. 83, 1075–1088. 10.1002/ana.25220.

8. Ouahchi, K., Arita, M., Kayden, H., Hentati, F., Hamida, M.B., Sokol, R., Arai, H., Inoue, K., Mandel, J.-L., and Koenig, M. (1995). Ataxia with isolated vitamin E deficiency is caused by mutations in the α–tocopherol transfer protein. Nat. Genet. 9, 141–145. 10.1038/ng0295-141.

9. Naureckiene, S., Sleat, David.E., Lackland, H., Fensom, A., Vanier, M.T., Wattiaux, R., Jadot, M., and Lobel, P. (2000). Identification of HE1 as the Second Gene of Niemann-Pick C Disease. Science 290, 2298–2301. 10.1126/science.290.5500.2298.

10. Carstea, E.D., Morris, J.A., Coleman, K.G., Loftus, S.K., Zhang, D., Cummings, C., Gu, J., Rosenfeld, M.A., Pavan, W.J., Krizman, D.B., et al. (1997). Niemann-Pick C1 Disease Gene: Homology to Mediators of Cholesterol Homeostasis. Science 277, 228–231. 10.1126/science.277.5323.228.

11. Fayngerts, S.A., Wu, J., Oxley, C.L., Liu, X., Vourekas, A., Cathopoulis, T., Wang, Z., Cui, J., Liu, S., Sun, H., et al. (2014). TIPE3 is the transfer protein of lipid second messengers that promote cancer. Cancer Cell 26, 465–478. 10.1016/j.ccr.2014.07.025.

12. Olayioye, M.A., Vehring, S., Müller, P., Herrmann, A., Schiller, J., Thiele, C., Lindeman, G.J., Visvader, J.E., and Pomorski, T. (2005). StarD10, a START domain protein overexpressed in breast cancer, functions as a phospholipid transfer protein. J. Biol. Chem. 280, 27436–27442. 10.1074/jbc.M413330200.

13. Vassilev, B., Sihto, H., Li, S., Hölttä-Vuori, M., Ilola, J., Lundin, J., Isola, J., Kellokumpu-Lehtinen, P.-L., Joensuu, H., and Ikonen, E. (2015). Elevated levels of StAR-related lipid transfer protein 3 alter cholesterol balance and adhesiveness of breast cancer cells: potential mechanisms contributing to progression of HER2-positive breast cancers. Am. J. Pathol. 185, 987–1000. 10.1016/j.ajpath.2014.12.018.

14. Heering, J., Weis, N., Holeiter, M., Neugart, F., Staebler, A., Fehm, T.N., Bischoff, A., Schiller, J., Duss, S., Schmid, S., et al. (2012). Loss of the ceramide transfer protein augments EGF receptor signaling in breast cancer. Cancer Res. 72, 2855–2866. 10.1158/0008-5472.CAN-11-3069.

15. Lee, A.J.X., Roylance, R., Sander, J., Gorman, P., Endesfelder, D., Kschischo, M., Jones, N.P., East, P., Nicke, B., Spassieva, S., et al. (2012). CERT depletion predicts chemotherapy benefit and mediates cytotoxic and polyploid-specific cancer cell death through autophagy induction. J. Pathol. 226, 482–494. 10.1002/path.2998.

16. Koga, Y., Ishikawa, S., Nakamura, T., Masuda, T., Nagai, Y., Takamori, H., Hirota, M., Kanemitsu, K., Baba, Y., and Baba, H. (2008). Oxysterol binding protein-related protein-5 is related to invasion and poor prognosis in pancreatic cancer. Cancer Sci. 99, 2387–2394. 10.1111/j.1349-7006.2008.00987.x.

17. Du, X., Zadoorian, A., Lukmantara, I.E., Qi, Y., Brown, A.J., and Yang, H. (2018). Oxysterol-binding protein-related protein 5 (ORP5) promotes cell proliferation by activation of mTORC1 signaling. J. Biol. Chem. 293, 3806–3818. 10.1074/jbc.RA117.001558.

18. Park, J.-S., and Neiman, A.M. (2020). XK is a partner for VPS13A: a molecular link between Chorea-Acanthocytosis and McLeod Syndrome. Mol. Biol. Cell 31, 2425–2436. 10.1091/mbc.E19-08-0439-T.

19. Kaplan, M.R., and Simoni, R.D. (1985). Intracellular transport of phosphatidylcholine to the plasma membrane. J. Cell Biol. 101, 441–445. 10.1083/jcb.101.2.441.

20. Kaplan, M.R., and Simoni, R.D. (1985). Transport of cholesterol from the endoplasmic reticulum to the plasma membrane. J. Cell Biol. 101, 446–453. 10.1083/jcb.101.2.446.

21. Newman, R.H., Fosbrink, M.D., and Zhang, J. (2011). Genetically Encodable Fluorescent Biosensors for Tracking Signaling Dynamics in Living Cells. Chem. Rev. 111, 3614–3666. 10.1021/cr100002u.

22. Wills, R.C., Goulden, B.D., and Hammond, G.R.V. (2018). Genetically encoded lipid biosensors. Mol. Biol. Cell 29, 1526–1532. 10.1091/mbc.E17-12-0738.

23. John Peter, A.T., Petrungaro, C., Peter, M., and Kornmann, B. (2022). METALIC reveals interorganelle lipid flux in live cells by enzymatic mass tagging. Nat. Cell Biol. 24, 996–1004. 10.1038/s41556-022-00917-9.

24. Hogan, P.G., and Rao, A. (2015). Store-operated calcium entry: Mechanisms and modulation. Biochem. Biophys. Res. Commun. 460, 40–49. 10.1016/j.bbrc.2015.02.110.

25. Chung, J., Torta, F., Masai, K., Lucast, L., Czapla, H., Tanner, L.B., Narayanaswamy, P., Wenk, M.R., Nakatsu, F., and De Camilli, P. (2015). PI4P/phosphatidylserine countertransport at ORP5-and ORP8-mediated ER–plasma membrane contacts. Science 349, 428–432. 10.1126/science.aab1370.

26. Sleight, R.G., and Pagano, R.E. (1983). Rapid appearance of newly synthesized phosphatidylethanolamine at the plasma membrane. J. Biol. Chem. 258, 9050–9058.

27. Baumann, N.A., Sullivan, D.P., Ohvo-Rekilä, H., Simonot, C., Pottekat, A., Klaassen, Z., Beh, C.T., and Menon, A.K. (2005). Transport of newly synthesized sterol to the sterol-enriched plasma membrane occurs via nonvesicular equilibration. Biochemistry 44, 5816–5826. 10.1021/bi048296z.

28. Stefan, C.J., Manford, A.G., Baird, D., Yamada-Hanff, J., Mao, Y., and Emr, S.D. (2011). Osh proteins regulate phosphoinositide metabolism at ER-plasma membrane contact sites. Cell 144, 389–401. 10.1016/j.cell.2010.12.034.

29. Saheki, Y., Bian, X., Schauder, C.M., Sawaki, Y., Surma, M.A., Klose, C., Pincet, F., Reinisch, K.M., and De Camilli, P. (2016). Control of plasma membrane lipid homeostasis by the extended synaptotagmins. Nat. Cell Biol. 18, 504–515. 10.1038/ncb3339.

30. Schauder, C.M., Wu, X., Saheki, Y., Narayanaswamy, P., Torta, F., Wenk, M.R., De Camilli, P., and Reinisch, K.M. (2014). Structure of a lipid-bound extended synaptotagmin indicates a role in lipid transfer. Nature 510, 552–555. 10.1038/nature13269.

31. Bumpus, T.W., and Baskin, J.M. (2016). A Chemoenzymatic Strategy for Imaging Cellular Phosphatidic Acid Synthesis. Angew. Chem. Int. Ed. 55, 13155–13158. 10.1002/anie.201607443.

32. Bumpus, T.W., and Baskin, J.M. (2017). Clickable Substrate Mimics Enable Imaging of Phospholipase D Activity. ACS Cent. Sci. 3, 1070–1077. 10.1021/acscentsci.7b00222.

33. Bumpus, T.W., Liang, F.J., and Baskin, J.M. (2018). Ex Uno Plura: Differential Labeling of Phospholipid Biosynthetic Pathways with a Single Bioorthogonal Alcohol. Biochemistry 57, 226–230. 10.1021/acs.biochem.7b01021.

34. Liang, D., Wu, K., Tei, R., Bumpus, T.W., Ye, J., and Baskin, J.M. (2019). A real-time click chemistry imaging approach reveals stimulus-specific subcellular locations of phospholipase D activity. Proc. Natl. Acad. Sci. U. S. A. 116, 15453–15462. 10.1073/pnas.1903949116.

35. Bumpus, T.W., Liang, D., and Baskin, J.M. (2020). IMPACT: Imaging phospholipase D activity with clickable alcohols via transphosphatidylation. Methods Enzymol. 641, 75–94. 10.1016/bs.mie.2020.04.037.

36. McDermott, M., Wakelam, M.J.O., and Morris, A.J. (2004). Phospholipase D. Biochem. Cell Biol. Biochim. Biol. Cell. 82, 225–253. 10.1139/o03-079.

37. Su, W., Yeku, O., Olepu, S., Genna, A., Park, J.-S., Ren, H., Du, G., Gelb, M.H., Morris, A. J., and Frohman, M.A. (2009). 5-Fluoro-2-indolyl des-chlorohalopemide (FIPI), a phospholipase D pharmacological inhibitor that alters cell spreading and inhibits chemotaxis. Mol. Pharmacol. 75, 437–446. 10.1124/mol.108.053298.

38. Min, S.-W., Chang, W.-P., and Südhof, T.C. (2007). E-Syts, a family of membranous Ca ^2+^ -sensor proteins with multiple C 2 domains. Proc. Natl. Acad. Sci. 104, 3823–3828. 10.1073/pnas.0611725104.

39. Bian, X., Saheki, Y., and De Camilli, P. (2018). Ca ^2+^ releases E-Syt1 autoinhibition to couple ER-plasma membrane tethering with lipid transport. EMBO J. 37, 219–234. 10.15252/embj.201797359.

40. Grimm, J.B., Muthusamy, A.K., Liang, Y., Brown, T.A., Lemon, W.C., Patel, R., Lu, R., Macklin, J.J., Keller, P.J., Ji, N., et al. (2017). A general method to fine-tune fluorophores for live-cell and in vivo imaging. Nat. Methods 14, 987–994. 10.1038/nmeth.4403.

41. Lambert, W.D., Scinto, S.L., Dmitrenko, O., Boyd, S.J., Magboo, R., Mehl, R.A., Chin, J.W., Fox, J.M., and Wallace, S. (2017). Computationally guided discovery of a reactive, hydrophilic trans-5-oxocene dienophile for bioorthogonal labeling. Org. Biomol. Chem. 15, 6640–6644. 10.1039/C7OB01707C.

42. Carlson, J.C.T., Meimetis, L.G., Hilderbrand, S.A., and Weissleder, R. (2013). BODIPY–Tetrazine Derivatives as Superbright Bioorthogonal Turn-on Probes. Angew. Chem. Int. Ed. 52, 6917–6920. 10.1002/anie.201301100.

43. Giordano, F., Saheki, Y., Idevall-Hagren, O., Colombo, S.F., Pirruccello, M., Milosevic, I., Gracheva, E.O., Bagriantsev, S.N., Borgese, N., and De Camilli, P. (2013). PI(4,5)P2-Dependent and Ca2+-Regulated ER-PM interactions mediated by the extended synaptotagmins. Cell 153, 1494. 10.1016/j.cell.2013.05.026.

44. Hollopeter, G., Lange, J.J., Zhang, Y., Vu, T.N., Gu, M., Ailion, M., Lambie, E.J., Slaughter, B. D., Unruh, J.R., Florens, L., et al. (2014). The membrane-associated proteins FCHo and SGIP are allosteric activators of the AP2 clathrin adaptor complex. eLife 3, e03648. 10.7554/eLife.03648.

45. Liang, D., Cheloha, R.W., Watanabe, T., Gardella, T.J., and Baskin, J.M. (2021). Activity-based, bioorthogonal imaging of phospholipase D reveals spatiotemporal dynamics of GPCR-Gq signaling. Cell Chem. Biol. 10.1016/j.chembiol.2021.05.020.

46. Tei, R., Bagde, S.R., Fromme, J.C., and Baskin, J.M. (2022). Activity-based directed evolution of a membrane editor in mammalian cells. Preprint at bioRxiv. 10.1101/2022.09.26.509516.

47. Alamudi, S.H., Satapathy, R., Kim, J., Su, D., Ren, H., Das, R., Hu, L., Alvarado-Martínez, E., Lee, J.Y., Hoppmann, C., et al. (2016). Development of background-free tame fluorescent probes for intracellular live cell imaging. Nat. Commun. 7, 11964. 10.1038/ncomms11964.

